# Systems-wide analysis of glycoprotein conformational changes by limited deglycosylation assay

**DOI:** 10.1101/2021.06.04.447131

**Authors:** Simon Ngao Mule, Livia Rosa-Fernandes, João V. P. Coutinho, Vinícius De Morais, Janaina Macedo da Silva, Verônica Feijoli Santiago, Daniel Quina, Gilberto Santos de Oliveira, Morten Thaysen-Andersen, Martin R. Larsen, Letícia Labriola, Giuseppe Palmisano

**Author notes:** To whom correspondence should be addressed: Prof. Giuseppe Palmisano, Glycoproteomics Laboratory Department of Parasitology, ICB, University of São Paulo, Brazil, Av. Prof. Lineu Prestes, 1374, 05508-900 - São Paulo – SP – Brazil.

## Abstract

A new method to probe the conformational changes of glycoproteins on a systems-wide scale, termed limited deglycosylation assay (LDA), is described. The method measures the differential rate of deglycosylation of N-glycans on natively folded proteins by the common peptide:N-glycosidase F (PNGase F) enzyme which in turn informs on their spatial presentation and solvent exposure on the protein surface hence ultimately the glycoprotein conformation. LDA involves 1) protein-level N-deglycosylation under native conditions, 2) trypsin digestion under denaturing conditions, 3) glycopeptide enrichment, 4) peptide-level N-deglycosylation and 5) quantitative MS-based analysis of the formerly N-glycosylated peptides. LDA was initially developed and the experimental conditions optimized using bovine RNase B and fetuin. The method was then applied to glycoprotein extracts from LLC-MK2 epithelial cells upon treatment with dithiothreitol to induce endoplasmic reticulum stress and promote protein misfolding. Data from the LDA and 3D structure analysis showed that glycoproteins predominantly undergo structural changes in loops/turns upon ER stress as exemplified with detailed analysis of ephrin-A5, GALNT10, PVR and BCAM. These results show that LDA accurately reports on systems-wide conformational changes of glycoproteins induced under controlled treatment regimes. Thus, LDA opens avenues to study glycoprotein structural changes in a range of other physiological and pathophysiological conditions relevant to acute and chronic diseases.

## Introduction

In the endoplasmic reticulum (ER), proteins are folded to their three-dimensional native conformational states depending on their amino acid sequence [1]. Proteins are folded by a complex molecular network composed of chaperones and protein folding enzymes regulated under physiological and stress conditions [2–4]. However, intracellular stress can impair ER quality control (ERQC) of proteins destined for secretion or transport to membranes, leading to the accumulation of misfolded and/or unfolded inactive precursor proteins in the ER causing ER stress [5–7]. Several diseases have been associated with a dysregulation of protein folding and accumulation of misfolded aggregates, including amyloidosis (e.g. Alzheimer’s disease) and prion diseases [8–10]. Protein kinase PKR-like ER kinase, inositol-requiring protein 1 alpha and activating transcription factor 6 alpha sense dysregulation of protein folding in the ER, leading to the activation of a network of signalling pathways termed the unfolded protein response (UPR), an evolutionary conserved mechanism serving to restore ER homeostasis and refold or degrade the unfolded proteins [11, 12]. UPR includes the downregulation of protein translation [13, 14], upregulation of protein-folding enzymes and chaperones [15], and upregulation of components of the ER-associated degradation (ERAD) pathway to enhance the degradation of the misfolded proteins [16, 17]. Proteins which cannot be refolded are targeted to the ERAD pathway, where they are ubiquitinated and degraded via the proteasomal pathway [18, 19]. In conditions of excess or sustained ER stress, autophagy can also be induced. If this process cannot revert the damage, it will end up triggering apoptosis culminating in cell death [20]. Chemicals such as tunicamycin (TM), dithiothreitol (DTT) and thapsigargin, which trigger ER stress through distinct mechanisms, have been used to study UPR [21, 22].

Newly synthesized proteins are co- and post-translationally modified with glycans in the ER followed by subsequent processing in the Golgi. This post-translational modification (PTM) is important for the correct folding and functioning of the proteins. Glycosylation is a vital and ubiquitous PTM, which, in eukaryotes, modifies over half of the proteome [23]. The *en bloc* transfer of pre-assembled oligosaccharides from a lipid-linked oligosaccharide donor to proteins in the ER at asparagine residues (Asn/N) in the conserved sequon (Asn(Xxx)Ser/Thr/Cys; where Xxx ≠ Pro) and the subsequent processing and maturation in the Golgi is the first step of the N-glycosylation pathway [24]. Protein glycosylation and the repertoire of glycans carried by newly synthesized glycoproteins is determined by the nature of the proteins undergoing glycosylation [25, 26] as well as a host of factors of the complex glycosylation machinery including the level of glycosylation enzymes [27], the availability of sugar nucleotide donors, subcellular localization [28], and cellular metabolism [29]. Glycans play diverse and important biological and structural functions including impacting the protein folding and localization, cell adhesion, function and regulation, stability of secreted proteins, protein solubility, glycan dependent regulation of receptor signalling and host-pathogen interaction [30–38]. The dysregulation of protein glycosylation has been associated with several human diseases including congenital disorders of glycosylation, diabetes mellitus, infectious diseases, auto-immune diseases, prion disease, cancer, cardiac rhythm disorders and chronic inflammatory diseases [39–41]. N-glycan dependent quality control of glycoproteins is regulated by lectin chaperones such as calnexin and calreticulin, disulphide isomerase, glucosyltransferases and glucosidases that prevent unfolded, misfolded and aggregated glycoproteins to exit the ER and enter the Golgi compartments. The presence of specific N-glycan structures within the ER correlate with the folding status and age of glycoproteins directing the activity of the folding machinery [42, 43].

X-ray crystallography and two-dimensional nuclear magnetic resonance (NMR) spectroscopy techniques are the major techniques applied to elucidate macromolecule structures including proteins [44, 45], and more recently, cryo-electron microscopy (cryoEM) [46]. These techniques, however, suffer technological setbacks including the need of highly purified protein crystals for X-ray crystallography, the need for high concentrations of non-aggregating proteins for NMR [47], and the need for sample structural homogeneity for cryoEM. The application of these techniques for glycoproteins is hindered by the size and heterogeneity of the oligosaccharide chains and the surface entropy of this PTM which impacts the crystallization. Limited proteolysis (LiP) is an alternative method for protein structure analysis of complex biological samples at scale [48]. LiP probes the conformation of proteins by employing peptidases which perform limited proteolysis on surface exposed peptide bonds of natively folded proteins [49]. The most recent development of LiP-MS is based on a dual-digestion using firstly a non-specific protease applied under native conditions followed by trypsin digestion under denaturing conditions. The resulting peptides are analysed by quantitative shotgun or targeted mass spectrometry enabling the identification of the structure-dependent proteolytic patterns [49]. This method has not explored the surface accessibility of glycoproteins.

We have here developed a new method for the large-scale analysis of glycoprotein conformational changes using limited deglycosylation assay (LDA). Previously it was shown that cells overexpressing endogenous PNGase F could be used as a sensor for studying the folding state of specific glycoproteins measured by differential SDS-PAGE migration [50, 51]. In the LDA method described herein, we have extended the concepts of limited proteolysis and deglycosylation sensitivity of differentially folded glycoproteins using large-scale differential and limited PNGase F deglycosylation.

## Materials and Methods

### Proof-of-concept and method optimization

Proof-of-concept and optimization of the LDA method were performed using two standard glycoproteins, bovine RNase B and fetuin (New England Biolabs) by PNGase F (New England Biolabs) under native conditions. The subsequent detection of the partially deglycosylated glycoproteins were performed by differential band migration on 15% SDS-PAGE. Detergent concentration (Nonidet P-40/NP-40) in the lysis buffer (adopted from [51]), PNGase F to glycoprotein ratio, incubation temperature and time were optimized. Final concentrations of 0.4% and 1% (v/v) NP-40 in the lysis buffer were tested. The optimal PNGase F (Cat. P0709S, NEB) concentration (relative to the glycoprotein substrate) was tested ranging from 1000, 500, 250, 125 to 75 U/ 5 μg RNase B (Cat.P7817S, NEB) and fetuin (Cat. P6042S, NEB). Different incubation times (2, 4, 8 and 16 h) were assayed, in addition to three temperature points (4°C, 24°C and 37°C). In all optimization conditions, fully deglycosylated and undegycosylated RNase B and fetuin were included as controls. The fully denatured standard glycoproteins were prepared as recommended by the manufacturer (NEBs). Briefly, 5 μg standard glycoprotein was mixed in 1 μL of 10× glycoprotein denaturing buffer (NEB) (final conc: 0.5% SDS, 40 mM DTT) and milli-Q ultrapure^®^ H_2_O to 10 μL final reaction volume and boiled for 10 min at 95°C, followed by sample cooling on ice. Subsequently, deglycosylation of the denatured glycoproteins was performed in a 20 μL reaction volume consisting of 10 μL of the previously denatured glycoprotein, 2 μL 10% NP-40 (final concentration 1% NP-40), 2 μL 10X GlycoBuffer 2 (final concentration 50 mM sodium phosphate (pH 7.5), 250 U PNGase F and 6 μL milli-Q ultrapure H_2_O. Deglycosylation was performed at 37°C for 1 h. Deglycosylation on the standard proteins was analyzed by gel electrophoresis on 15% SDS-PAGE gels and the gels stained by Coomassie Brilliant Blue R-250 Dye (Cat. 20278, Thermo Scientific). The optimal conditions were applied to study limited deglycosylation on LLC-MK2 cellular glycoproteins upon induction of ER stress, as described below.

### Cell cultures

Rhesus monkey kidney epithelial cells (LLC-MK2) (ATCC, USA) [52] were maintained in RPMI media (Life Technologies) supplemented with 10% (v/v) heat-inactivated fetal bovine serum (FBS) and 100 U/mL penicillin in 75 cm^2^ culture flasks at 37°C with 5% CO_2_ until the cells reached 90% confluence. Passaging was done by seeding 1.0 × 10^6^ cells in new 75 cm^2^ culture flasks. Cell cultures were periodically assayed for mycoplasma contamination using PCR test.

### ER stress induction, cell lysis under native conditions and protein extraction

LLC-MK2 cells were seeded and grown in RPMI media supplemented with 10% FBS at 37°C with 5% CO_2_ to 90% confluence. The cells were subsequently rinsed 3 times with 1× PBS (137 mM NaCl, 2.7 mM KCl, 10 mM Na_2_HPO_4_, 1.8 mM KH_2_PO_4_, pH 7.4), treated with 3 mM dithiothreitol (DTT) (molecular grade, Cat. 3483-12-3, Promega), and incubated for 30 min at 37°C in 5% CO_2_ in serum-free RPMI media. This concentration of DTT has previously been shown to induce ER stress in several cell lines [53–55]. Control conditions of LLC-MK2 cells incubated in the absence of DTT were included. After incubation, the cells were washed 3 times with ice-cold 1 × PBS.

Subsequently, the cells were detached by cell scraping technique in 2 mL ice-cold 1× PBS, and transferred into pre-chilled Protein LoBind® 2 mL microtubes (Eppendorf AG, Hamburg, Germany). The cells were pelleted by low-speed centrifugation at 800×g for 3 min at 4°C. The supernatant was discarded by aspiration, and the cells were resuspended in 200 μL ice-cold cell lysis buffer adopted from [50, 51] with minor modifications (50 mM Tris/HCl (pH 8.0), 0.4% NP-40, and 150 mM NaCl) containing protease inhibitor cocktail (1×) (Promega), followed by cell lysis by incubation on ice for 20 min. Ultimately, the lysed cell debris were cleared by centrifugation at 17,800×g for 10 min at 4°C and the supernatants transferred separately into new pre-chilled Protein LoBind® microtubes placed on ice. The samples were submitted to total proteome analysis and LDA. Proteins were assayed using SDS-PAGE, and the proteome from both control and ER stress-induced fractions submitted to sample processing prior to label free nLC-MS/MS analysis as described below.

To determine the protein concentration, cells were cultured as previously described and resuspended in 8 M urea supplemented with 1X protease inhibitors, lysed by 3 freeze thaw cycles, cleared by centrifugation as before, and the proteins quantified by Qubit fluorometric detection method.

### Optimized LDA on cellular proteins

The optimized conditions previously determined were applied to LLC-MK2 cellular glycoproteins and confirmed by lectin blot analysis prior to nLC-MS/MS analysis. These conditions were: PNGase F (1000 U/100 μg protein), 16 h incubation at 4°C in 200 μL lysis buffer with 0.4% NP-40. Biological replicates for each condition with and without PNGase F treatment were analyzed. PNGase F was deactivated by boiling the samples for 10 min at 95°C in Laemmli sample buffer (final concentration of 31.5 mM Tris, pH 6.8, 10% glycerol, 1% SDS, 10 mM DTT, 0.005% bromophenol blue) [56].

### SDS-PAGE and lectin blotting

To visualize the effect of DTT on (glyco)protein expression, 10 μg protein extract from LLC-MK2 cells treated with or without 3 mM DTT was separated in 12% SDS-PAGE at a constant voltage of 125 V, stained by CBB, destained and the protein profiles acquired using a ChemiDoc Imaging system (Bio-Rad). Limited deglycosylation of LLC-MK2 cellular glycoproteins treated with or without 3 mM DTT, and treated with or without PNGase F was assayed by lectin blotting using Concanavalin A (Con A) (Cat. B-1005-5, Vector Laboratories). A total of 10 μg protein each were electrophoresed in a 12% SDS-PAGE gel and transferred onto PVDF membranes at a constant of 300 mA for 2 h. The membranes were blocked with 5% BSA and probed overnight at 4°C with biotinylated Con A lectin followed by horseradish peroxidase (HRP) secondary antibody for 1 h at RT with orbital shaking. The lectin blots were developed using SuperSignal™ West Pico Plus Chemiluminescent substrate (Thermo Fischer Scientific) and the images acquired using a ChemiDoc Imaging system (Bio-Rad). The band intensities were analyzed using the ImageLab (BioRad) software. Statistical analyses of the lectin intensities were calculated in GraphPad (v.8) using one-way ANOVA followed by Tukey's post-hoc test, with a p < 0.05 considered statistically significant.

### Sample processing, protein digestion and peptide extraction

To facilitate removal of NP-40 detergent prior to LC-MS/MS analysis, we performed in-gel protein digestion and peptide extraction [57–59]. Briefly, proteins were introduced into 12% SDS-PAGE gels using a constant voltage (125 V) to approximately 1 cm into the resolving gel. The proteins were stained by CBB, destained in 30% methanol, 10% glacial acetic acid and 60% distilled H_2_O, followed by excision of stained gel bands. The gel bands were completely destained, treated with 10 mM DTT at 56°C for 45 min, 55 mM IAA at room temperature for 30 min in the dark and digested at 37°C for 16 h with 2 μg porcine trypsin (sequencing grade, Promega). The resultant tryptic peptides were extracted in 40% ACN/0.1% TFA into fresh Protein LoBind® microtubes, dried by vacuum centrifugation, and resuspended in 50 μL 0.1% TFA. The peptides were desalted using oligo™ R3 reversed-phase resin (Cat. # 1133903, Thermo Fisher) and subsequently dried as before.

### Glycopeptide enrichment and deglycosylation

Hydrophilic interaction liquid chromatography (HILIC) was employed to enrich for glycopeptides [60–65]. Briefly, the desalted tryptic peptides were passed three times through a p200 pipette tip packed onto a C8 disk (Empore) with PolyHYDROXYETHYL ATM HILIC SPE resin (PolyLC Inc). The microcolumns were washed using 80% (v/v) ACN/1% (v/v) TFA, followed by the elution of the bound glycopeptides in 50 μL 0.1% (v/v) TFA, 50 μL 25 mM NH_4_HCO_3_ and finally in 50 μL 50% (v/v) ACN. The three fractions were combined, dried by vacuum centrifugation, resuspended in 20 μL 50 mM NH_4_HCO_3_ and PNGase F (250 U) added. Deglycosylation was performed for 16 h at 37°C prior to desalting on two C18 disks (3M Empore C18 disks, product number 2215) packed in a p200 tip [66].

### nLC-MS/MS analysis

Formerly N-glycopeptides were separated by nano-LC-MS/MS coupled to an ESI-LTQ-Orbitrap Velos mass spectrometer (Thermo Fisher Scientific). Mass spectrometric analysis was performed with the following settings: The injection volume was 10 μL, and the total analysis time for each sample was 105 min. The 20 most intense precursors selected from the Orbitrap MS1 full scan (resolving power 120,000 full width at half-maximum (FWHM) @ m/z 200) were isolated and fragmented by collision-induced dissociation (CID) and detected in the dual-pressure linear ion trap with 35 as normalized collision energy (NCE). The dynamic exclusion duration was set to 15 s.

Identification and quantification of proteins and peptides in the formerly N-glycosylated peptide (FNGPs) fraction were performed using MaxQuant v.1.5.3.8 [67] using the label-free quantification (LFQ) algorithm. The MS/MS spectra were searched against the *Macaca mulatta* proteome database in UniProtKB (downloaded, April 2020 with 44,402 entries). Trypsin was set as the proteolytic enzyme with 2 missed cleavages, and the minimum peptide length was set at 7 amino acid residues. The precursor ion and fragment ion mass tolerance were set at 20 ppm and 0.5 Da, respectively while the protein FDR was set at 0.01. Carbamidomethylation of cysteines was set as a fixed modification, while deamidation (N) was set as a variable modification and used for protein quantification. Localization probability was calculated using the algorithm embedded in MaxQuant [68].

### Quantitative mass spectrometry analysis of the total proteome

Proteome-level changes in LLC-MK2 cells treated with or without DTT were evaluated. The total proteins were extracted as previously described, followed by in-gel denaturation and alkylation steps using 10 mM DTT and 55 mM IAA, respectively. The proteins were in gel-digested overnight with sequencing grade trypsin (Promega). The resulting tryptic peptides were desalted using in-house C18 disks, and the peptides submitted to nLC-MS/MS analysis as previously described. Carbamidomethylation of cysteines was set as a fixed modification, while oxidation of methionine and protein N-terminal acetylation were set as the variable modifications, and used for protein quantification. nLC-MS/MS analysis was performed as previously described.

### Bioinformatics and data analysis

Statistical analyses were performed in Perseus v.1.6.10.43 [69] and *GraphPad Prism* v.8.0 (*GraphPad* Software, La Jolla California USA, www.graphpad.com). Contaminants and proteins identified in the reverse database were excluded before statistical analyses. For both total proteome and deglycoproteome analyses, a cut-off was included to filter for proteins and FNGPs, respectively, which were identified and quantified in duplicates in at least one condition. Regulated proteins between LLC-MK2 cells treated with and without DTT were analyzed by student t-test using Benjamini-Hochberg based FDR (FDR < 0.05) to test the effect of DTT on protein expression.

Analysis of FNGPs was performed to evaluate the effect of DTT on 1) N-glycosylation and 2) structural conformational changes, respectively, as described below. Deamidated peptides with the glycosylation motif ((Asn[Xxx]Ser/Thr/Cys) where Xxx ≠ Pro))[70] and localization probabilities ≥ 0.9 (90%) were considered for subsequent analyses. t-test analysis of LFQ values of the protein groups from control and DTT treated cells in the absence of PNGase F treatment was performed using Benjamini-Hochberg based FDR (FDR < 0.05). The total ion intensities of peptides from control cells (treated with or without PNGase F) and DTT treated cells (with or without PNGase F) were evaluated using ANOVA with Tukey’s test to correct for multiple comparisons in *GraphPad* Prism v.8.0. Statistical analysis of FNGPs was performed by ANOVA using Benjamini-Hochberg FDR (FDR < 0.05).

### Gene ontology analysis

Gene ontology (GO) analysis of the glycoproteins identified and quantified by mass spectrometry analysis was determined using the g:profiler tool [71]. Enriched cellular components, molecular functions and biological processes were determined by applying a q-value cut-off of 0.05, corrected by Benjamini-Hochberg FDR.

### Glycoprotein structure analysis

Selected glycoproteins with pronounced structural changes upon DTT treatment were analyzed to determine their 3D structures and localization of the N-glycosylation sites based on available crystal structures of homologous proteins deposited in the Protein Data Bank (PDB) (https://www.rcsb.org/). For this analysis, the Protein Homology/analogY Recognition Engine v.2.0 (Phyre^2^) [72] was used to model and predict the 3D structure of selected glycoproteins identified by mass spectrometry displaying pronounced structural changes upon DTT treatment. The glycoprotein 3D structures were analyzed for their localization of the mapped N-glycosites and solvent accessible surface area (SASA) of the modified N-glycosylation sites using Pymol v.2.3.4, with dot density set to 4, and a solvent radius of 1.4 Å. To enable 3D structure alignment and comparison with homologous glycoprotein structures already deposited in the PDB, PyMod plugin of PyMOL [73–75] was used for alignment to determine if the positions undergoing pronounced structural modulations upon DTT treatment were conserved between *M. mulutta* and *H. sapiens* and/or *M. musculus* orthologous glycoproteins. For this, previously modelled LLC-MK2 glycoproteins from our study were loaded on PyMod, and a PDB database search using remote BLAST performed using a E-value threshold of 10 was used to retrieve similar 3D-structures in PDB. High scoring PDB structures were downloaded and the 3D structures aligned. The N-glycosites undergoing pronounced conformational changes based on our analysis were mapped on the homologous glycoproteins and illustrated.

In addition, identified N-glycosites undergoing pronounced and subtle structural changes upon DTT treatment were analyzed using Prediction of Order and Disorder by evaluation of NMR data (ODiNPred) [76], a sequence-based protein order/disorder predictor. The evolution option was activated for this analysis.

## Results and Discussion

### Proof-of-concept and optimization of the limited deglycosylation assay (LDA)

In this study, we introduce the limited deglycosylation assay (LDA) to study conformational changes of glycoproteins on a systems-wide scale using a quantitative label-free analysis of deglycosylated peptides. The experimental workflow for LDA is summarized in **Figure 1**. In step 1, standard glycoproteins, cells or tissues are subjected to biotic or abiotic stress influencing glycoprotein folding and structure. Proteins are extracted under native conditions using a mild detergent to improve membrane solubilization while avoiding disruption of protein conformation (step 2), before treatment of the still natively folded proteins with PNGase F (step 3). The glycoprotein folding states dictate the susceptibility to PNGase F-mediated removal of their surface exposed N-glycans. Subsequently, proteins are loaded into a SDS-PAGE to remove the detergent and subsequently in-gel digested using trypsin (step 4). Glycosylated peptides are enriched using HILIC (step 5) and subjected to deglycosylation using PNGase F (step 6). Resulting FNGPs are analysed using label-free quantitative mass spectrometry (MS)-based proteomics. The differential ion intensity of FNGPs allow a site-specific mapping of PNGase F accessibility which can be correlated with conformational changes in the 3D structures upon stress.

**Figure 1.**
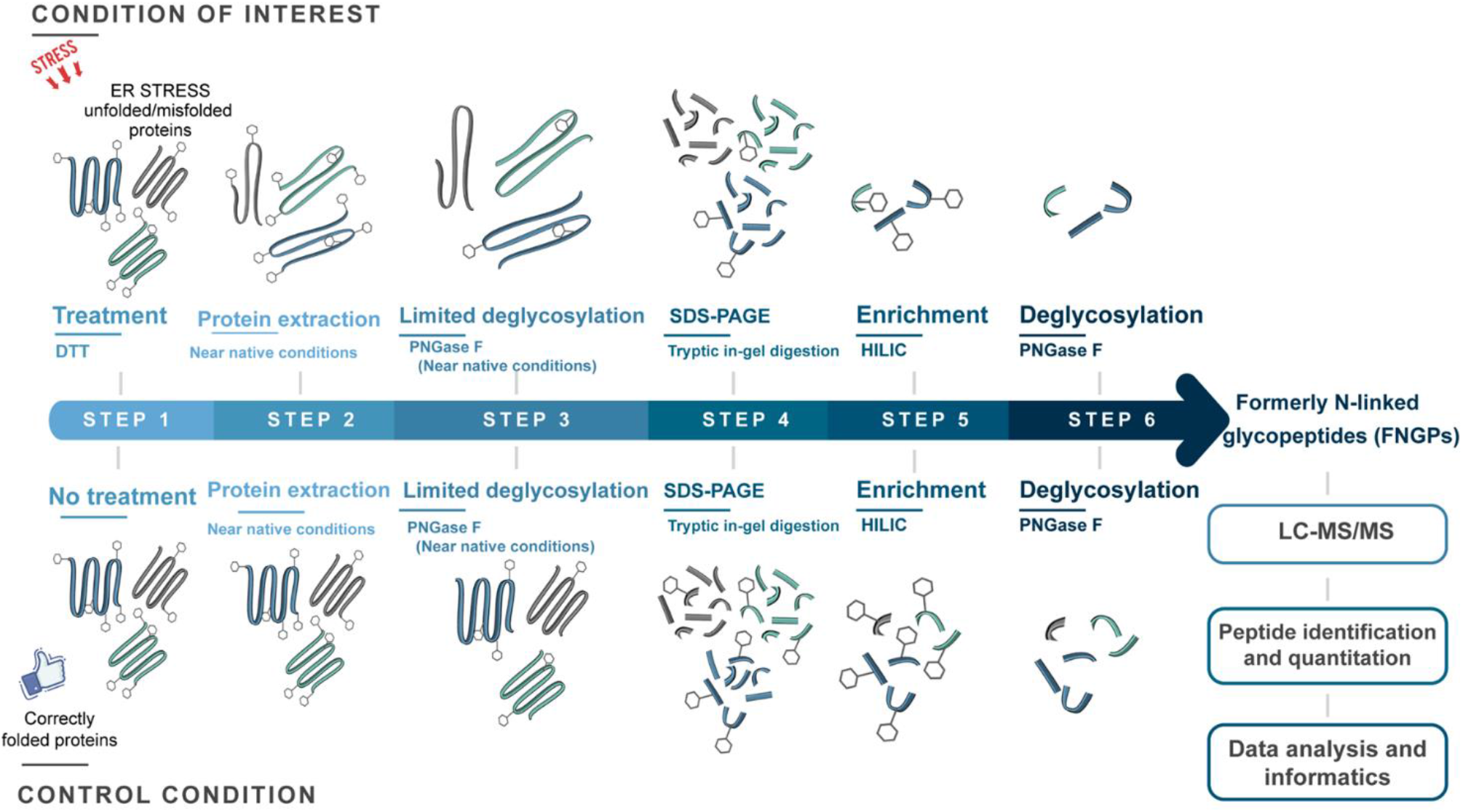
Experimental workflow of the limited deglycosylation assay (LDA). LDA probes glycoprotein conformational changes on a global scale following perturbation-induced structural modulation. Cells treated or not with ER stress inducers are lysed in native conditions and subjected to PNGase F treatment (4°C for 16 h). The resulting proteins are analyzed by GeLC approach using SDS-PAGE and in-gel digestion of the entire lane. The tryptic glycopeptides are enriched using HILIC SPE and subsequently deglycosylated by overnight PNGase F treatment. Quantitative nLC-MS/MS analysis was performed to identify differentially formerly N-glycopeptides (FNGPs).

Initially, the LDA method was optimized using RNase B and Fetuin to establish the optimal detergent concentration, PNGase F-to-glycoprotein ratio, and the reaction time and temperature (**Figure 2**). Bovine RNase B carries high mannose-type glycans at a single N-glycosite, while fetuin has 3 N- [61, 77–79] and 6 O-glycosites [80, 81]. The two glycoproteins were partially deglycosylated under native conditions, while their denatured forms, included as controls, were completely deglycosylated, as illustrated by their differential migration on SDS-PAGE. In our analysis, PNGase F hydrolysed N-glycans more efficiently in 0.4 % NP-40 compared to 1% NP-40 buffer for both RNase B and fetuin, as shown by the intensity of bands corresponding to the completely deglycosylated controls (**Figure 2A**). The impact of the deglycosylation temperature was evaluated at 4°C, 24°C and 37°C in native buffer with 0.4% NP-40, which showed a higher deglycosylation efficiency at higher temperatures (**Figure 2B**). At 37°C, the band corresponding to the fully deglycosylated band was observed, highlighting temperature-dependent changes in glycoprotein structural conformation and subsequent differential sensitivity of PNGase F at 37°C compared to 4°C and 24°C. Higher degree of deglycosylation of RNase B and fetuin at PNGase F:glycoprotein substrate ratio of 1000 U/2.5 μg (corresponding to 400 U/μg) at 4°C for 16 h in 0.4% NP-40 buffer was demonstrated, while decreasing rates of deglycosylation were demonstrated up to 15 U/μg (**Supplementary Figure 2C, 2D**). Incubation time of 16 h in native conditions demonstrated higher deglycosylation levels for both glycoproteins (**Figure 2E, F**). Taken together, native PNGase F-mediated deglycosylation was shown to be most efficient in 0.4 % NP-40 buffer for 16 h incubation at 4°C preserved the native structural forms of the glycoproteins. Thus, these conditions were used for the subsequent LDA experiments with cellular proteins following DTT-based induction of ER stress. PNGase F has low activity on native glycoproteins [82], while misfolded and/or unfolded/denatured glycoproteins are susceptible to cleavage of N-glycans by endogenous endoglycosidases [83, 84]. Endo H and PNGase A have even less activity on native glycoproteins [83]. Methods to probe the structural conformational states of standard glycoproteins using PNGase F coupled to SDS-PAGE, lectin blotting, mass spectrometry, NMR procedures have previously been demonstrated [82–84]. However, LDA is the first method to study the systems-wide glycoprotein structural modulation in a biological system.

**Figure 2.**
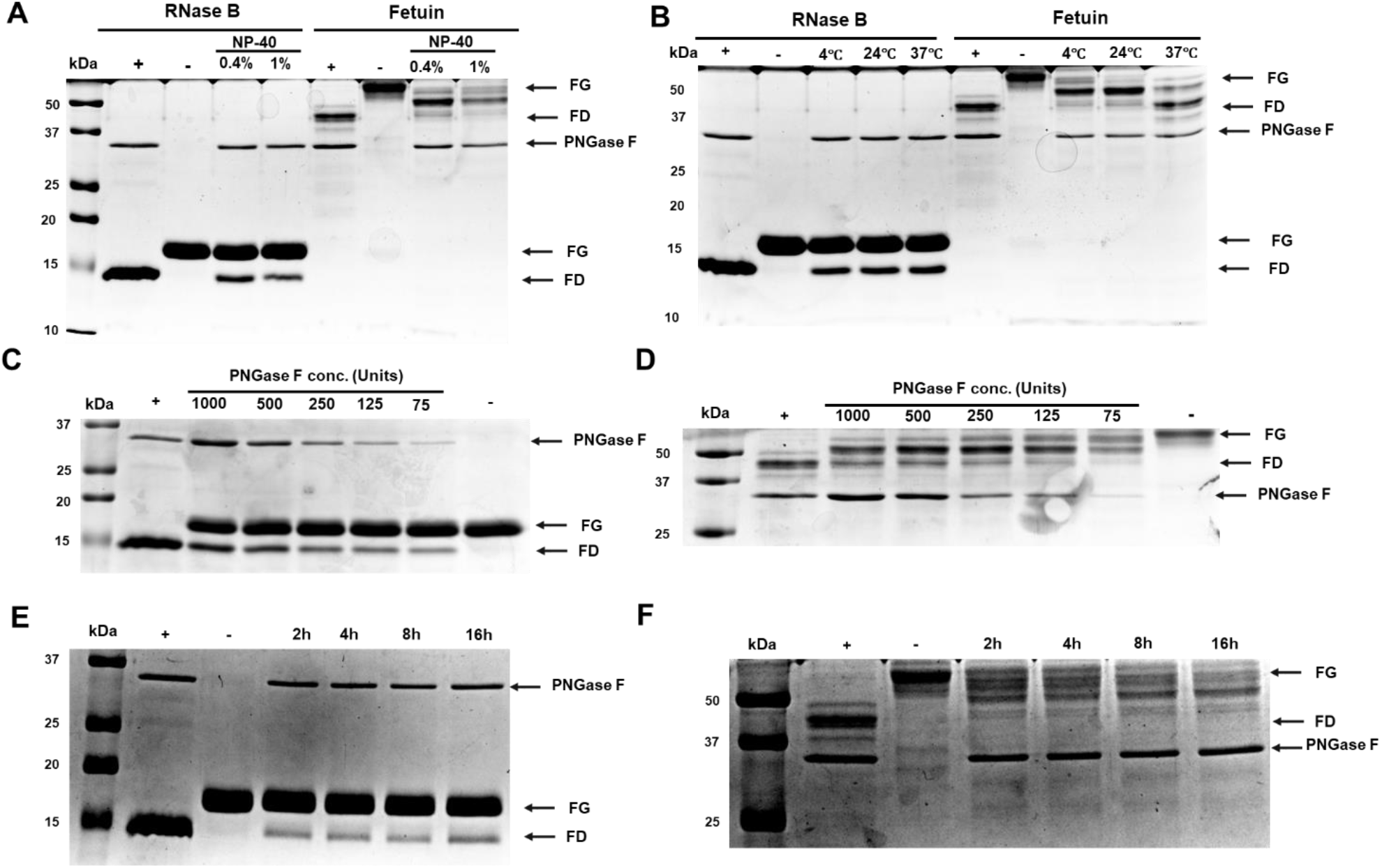
Optimization of the LDA method using RNase B and fetuin. Experimental conditions for optimal limited deglycosylation of bovine RNase B and fetuin by PNGase F were determined. The optimal **A**) NP-40 concentration, **B**) temperature, **C**) and **D**) PNGase F:glycoprotein ratio for RNase B and fetuin, respectively, and, **E**) and **F**) the optimal time for the deglycosylation of RNase B and fetuin, respectively, were assayed by differential migration of the glycoproteins in 15% SDS-PAGE. Fully deglycosylated (+) and fully glycosylated (−) proteins were included for each optimization reaction, denoted by FD and FG, respectively.

**Figure 3.**
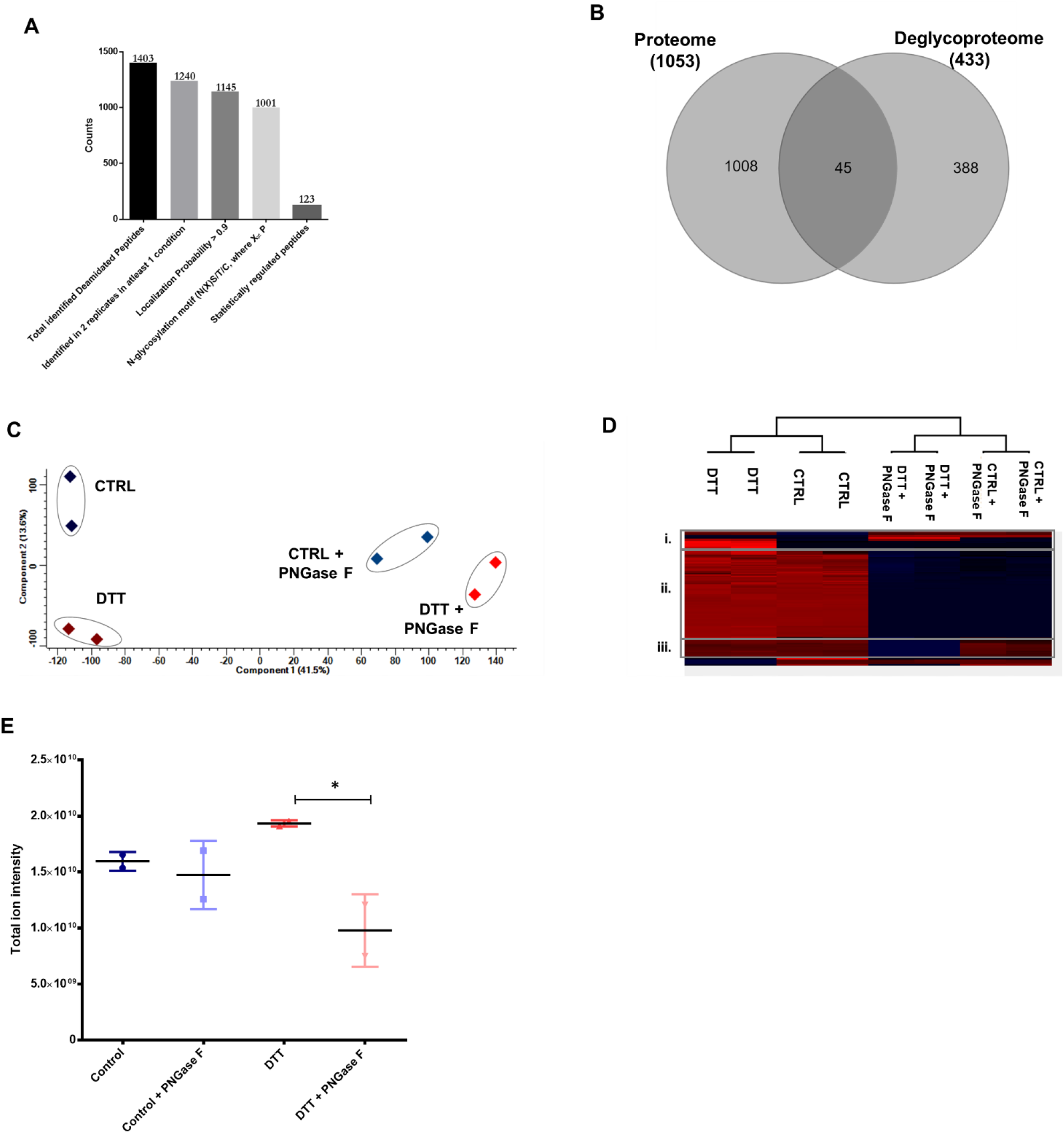
Limited deglycosylation of LLC-MK2 cells treated with or without 3 mM DTT. The mass spectrometry analysis showing **A**) number of identified and quantified deaminated peptides, **B**) Venn diagram showing shared proteins from proteome and deglycoproteome analysis, **C**) the PCA **D**) heatmap of deamidated peptides, **E**) total ion intensities of the deamidated deglycopeptides from control, control + PNGase F, DTT, and DTT + PNGase F fractions.

Proteins extracted from LLC-MK2 cells treated with or without 3 mM DTT were subjected to limited deglycosylation under native conditions and analyzed using lectin blotting to confirm the optimal conditions previously determined on standard glycoproteins (**Supplementary Figure 1A**). Cells were lysed in native lysis buffer with 0.4% NP-40, and treated overnight with 1000 U PNGase F at 4°C or left untreated. A denatured protein control was included, where equal amount of proteins was initially boiled in glycoprotein denaturing buffer for 10 min at 95°C, cooled on ice prior to overnight incubation with PNGase F as described above. Notably, the complete deglycosylation (0.09) of the denatured fraction was achieved analysing the ratio between lectin blot intensities in the +PNGase F divided by −PNGase F conditions (+PNGase F/−PNGase F = 0.09) (**Supplementary Figure 1B**). Under native conditions, however, limited deglycosylation was observed, where more deglycosylation of proteins from cells treated with 3 mM DTT (+PNGase F/−PNGase F = 0.39) compared to proteins from cells incubated in the absence of DTT (+PNGase F/−PNGase F = 0.59) (**Supplementary Figure 1B**).

These data support that PNGase F is able to differentially remove N-glycans from proteins with different structural conformations depending on its accessibility to the N-glycans on the natively folded glycoproteins. As shown for the denaturing conditions where the proteins are completely un- or misfolded (**Supplementary Figure 1**), all the N-glycans are hydrolyzed by PNGase F, illustrating their improved susceptibility to hydrolysis by the enzyme compared to the partial hydrolysis under native conformations. LDA on DTT treated cells revealed a higher deglycosylation rate compared to control cells grown in the absence of DTT, revealing conformation-based differential accessibility of PNGase F to N-glycans (**Supplementary Figure 1B**). Incubation of cells in DTT treated culture medium prevents the formation of disulphide bond in nascent polypeptide chains in the ER as well as reduction of disulphide bonds in already folded proteins, which results in conformational modulation and subsequent accumulation of misfolded/unfolded proteins in the ER [85, 86].

Proteome-wide analysis of LLC-MK2 cells treated with or without 3 mM DTT was performed prior to LDA to evaluate the proteome-wide modulation upon incubation with DTT. Protein band patterns were initially visualized by SDS-PAGE (**Supplementary Figure 2A**), followed by an in-depth bottom-up label-free mass spectrometry-based proteomics approach. No notable differences between control and DTT treated cells were detected through SDS-PAGE, as indicated by the protein bands intensity ratios between DTT and CTRL determined to be 1.06. Quantitative proteome data showed a separation of the control and DTT treated fractions by principal component analysis (PCA) (**Supplementary Figure 2B**). However, paired t-test analysis using Benjamini-Hochberg based FDR at an FDR < 0.05 showed that upon 3 mM DTT treatment for 30 min, the regulation in protein abundance at the protein level was not statistically significant (**Supplementary Figure 2C**). Longer incubation periods of 18 h with 2 mM DTT have been shown to cause clear UPR activation in Hela cells along with thapsigargin and tunicamycin [22]. In this study, LLC-MK2 cells were incubated for 30 min in the presence of 3 mM DTT, a short time to observe complete UPR activation, but sufficient time to analyze glycoprotein conformation changes prior to protein level abundance changes due to DTT induced ER stress.

A total of 1540 proteins were identified and quantified in this study using label-free mass spectrometry (**Supplementary Table 1**). Of these, 1053 proteins were identified in 2 replicates in at least one condition, and they were used for subsequent analyses. A total of 9 and 8 proteins were identified exclusively in control and DTT fractions, respectively (**Supplementary Table 1**). Analysis of the total quantitative proteome data revealed subtle increase in ER stress associated proteins such as the chaperones HSP90 β1 protein (I0FUR4/endoplasmin) and 78 kDa glucose-regulated protein (F7C3R1/HSPA5), calnexin (H9ES13/CANX), calreticulin (G7NLC3/CALR), and protein disulphide isomerases (A0A5F8ABB1/PDIA4; F6W4X0/PDIA3; and A0A1D5RKD4/P4HB) (**Supplementary Table 1**), suggesting the early onset of UPR.

LDA was performed using the optimized conditions and the deglycosylation rate was monitored using Con A blotting before trypsin digestion, glycopeptide enrichment and mass spectrometry analysis. Lectin blot intensities corroborated that more N-glycans were cleaved off in the DTT treated fractions compared to control fractions (**Supplementary Figure 3A)**. A statistically significant reduction in levels of glycosylation between DTT+PNGase F and DTT only fractions (ratio = 0.55) was seen using one-way ANOVA using Tukeys correction of multiple comparisons (p < 0.05) (**Supplementary Figure 3B**). In addition, higher levels of glycosylation were observed in DTT treated fractions compared to control fractions in the absence of PNGase F treatment (**Supplementary Figure 3B**). It has been shown that the prevention of disulphide bond formation and subsequent protein misfolding by treatment of cells with mild levels of DTT leads to complete glycosylation of N-sequons that, in native conformations, are missed or undergo variable levels of glycosylation [87, 88].

Analysis of the FNGPs by MS/MS following limited deglycosylation and HILIC enrichment identified a total of 1403 deamidated peptides, of which 1240 were identified in 2 replicates in at least one condition, representing 571 unique source glycoproteins (**Supplementary Table 2; Figure 2A**). To ensure high confidence of the deamidated peptides identified and quantified in this study, peptides with a localization probability ≥ 0.9 (90 %) were selected, resulting in 1145 deglycopeptides. Of these, 1001 peptides, corresponding to 433 unique source glycoproteins, had one or more N-glycosylation sequence motifs, and were considered for further analysis. Gene ontology analysis of the identified glycoproteins is illustrated in **Supplementary Figure 4**. Plasma membrane, organelle membrane and extracellular membrane-bounded organelles terms were enriched as cellular components (**Supplementary Fig. 4A**). Indeed, 93.7% of the total identified glycoproteins were associated to membrane showing the robustness of the N-linked glycopeptide enrichment and site analysis (**Supplementary Figure 4B**). Binding of cell adhesion molecules, transmembrane transporters and glycosyltransferases activity were enriched as molecular functions (**Supplementary Figure 4C**). The identified glycoproteins were associated to cell adhesion, integrin-mediated cell signaling, glycosylation, secretion and vesicle-mediated transport as biological processes (**Supplementary Figure 4D**).

Statistical analysis by ANOVA at an FDR < 0.05 showed that 123 peptides were statistically regulated between the four fractions (**Figure 2A**), representing 101 unique source glycoproteins. The enrichment step using HILIC in our current study is paramount to increase the coverage of glycopeptides, as revealed by Venn diagram analysis showing that only 45 glycoproteins were identified and quantified in common between the proteome and deglycoproteome datasets (**Figure 2B**). It should also be noted that measuring protein levels between the two conditions is also important to exclude any protein expression-dependent changes that can be confounding factors for determining differential abundance of FNGPs. In this study, a short stimulation with DTT did not influence the protein levels (**Supplementary Figure 2C**).

Deglycoproteome analysis by PCA and Euclidean distance showed that presence and absence of PNGase F treatment during the limited deglycosylation step 3 (**Figure 1**) accounted for the majority of the variation in the dataset (**Figure 2C and D**). A clear separation into 2 major clusters was observed based on the first principal component, which accounted for 41.5% of the total variation in the dataset (**Figure 2C**). Separation between DTT treated and control fractions in the absence of PNGase F treatment was observed based on the second principal component, representing 13.6% of total variation in the dataset, which highlighted the potential effect of DTT on LLC-MK2 cellular glycosylation (**Figure 2C**). Statistical analysis using ANOVA at an FDR < 0.05 identified 123 regulated FNGPs. The regulated FNGPs formed three major sub-clusters which included: i) glycopeptides whose abundances increased following DTT incubation compared to the control fractions in the absence of PNGase F treatment, and subsequently were completely deglycosylated in DTT fractions compared to their PNGase F untreated counterparts; ii) glycopeptides whose structural conformations in native and mis/unfolded conformations had equal accessibility by PNGase F, and iii) glycopeptides whose PNGase F accessibility was more following DTT incubation compared to the –DTT control condition. The structure of selected proteins in subset iii) was further analyzed and discussed.

Total ion intensities of FNGPs with the N-glycosylation sequence motif revealed statistically significant differential deglycosylation by PNGase F, showing that more N-glycans were accessible to PNGase F cleavage in the DTT treated fractions compared to native conditions (**Figure 2E**).

A total of 43 deamidated peptides were identified and quantified in subset iii), which included FNGPs missing exclusively in the DTT treated fractions incubated overnight in the presence of PNGase F. The corresponding glycoproteins were considered to have undergone pronounced structural modulation upon DTT treatment, exposing the N-glycans to PNGase F, leading to their complete cleavage (**Supplementary Table 2**). Three glycoproteins undergoing pronounced and/or subtle conformational modulation upon DTT treatment, with 1, 2 and 4 mapped N-glycosites were chosen for further discussion following 3D structural analysis.

One N-glycosite (N37) of Ephrin-A5 (F7GZC7/EFNA5), a membrane GPI-anchored protein, was identified with profound structural modulation. Analysis of the global breast cancer xenograft tissue proteome identified the human homolog to this protein to be glycosylated [89], with Hex_5-6_ HexNA_c2_ glycans at N37 reported in Glyconect. The human homolog ephrin-A5 ectodomain crystal structure has been determined by X-ray crystallography [90]. Using Phyre2, the 3D structure of the identified LLC-MK2 cellular ephrin-A5 was modelled, revealing that N37 is located on a loop, 3 amino acid residues away from a beta sheet (**Figure 4A**). Under native conditions, this site is completely shielded from PNGase F accessibility, as the ratio between CTRL + PNGase F and CTRL is 1.05. The SASA of ephrin-A5 N37 was determined to be 34.3 Å using PyMol, indicating that this site has a low solvent-exposed surface area (**Figure 4B**). Structure alignments of human ephrin-A5 (4LOP chain B) [91] and *Mus musculus* ephrinA5 (1SW chain A) [92] showed the conservation of N37 on a loop across species (**Figure 4C**), suggesting possible structural modulation of this protein in different organisms upon ER stress. This glycosite is shared within ephrin-A ligands. It has been shown that ephrin-A1 ligand glycosylation interferes with ephrin-A2 receptor binding, internalization and degradation affecting the downstream signalling pathways. The deglycosylated form of ephrin-A1 does not inhibit cancer cell migration and impairs ligand secretion showing the importance of ephrin-A ligands glycosylation [93]. Analysis of ordered and disordered localization of the N-glycosite using ODiNPred showed that at position N37, the probability of the N-glycosite to be located in a disordered region was 0.0342 (**Figure 4D**). All N37 neighbouring amino acid residues had low probabilities of disorder, in agreement with findings on N-glycosites of human proteins [94].

**Figure 4.**
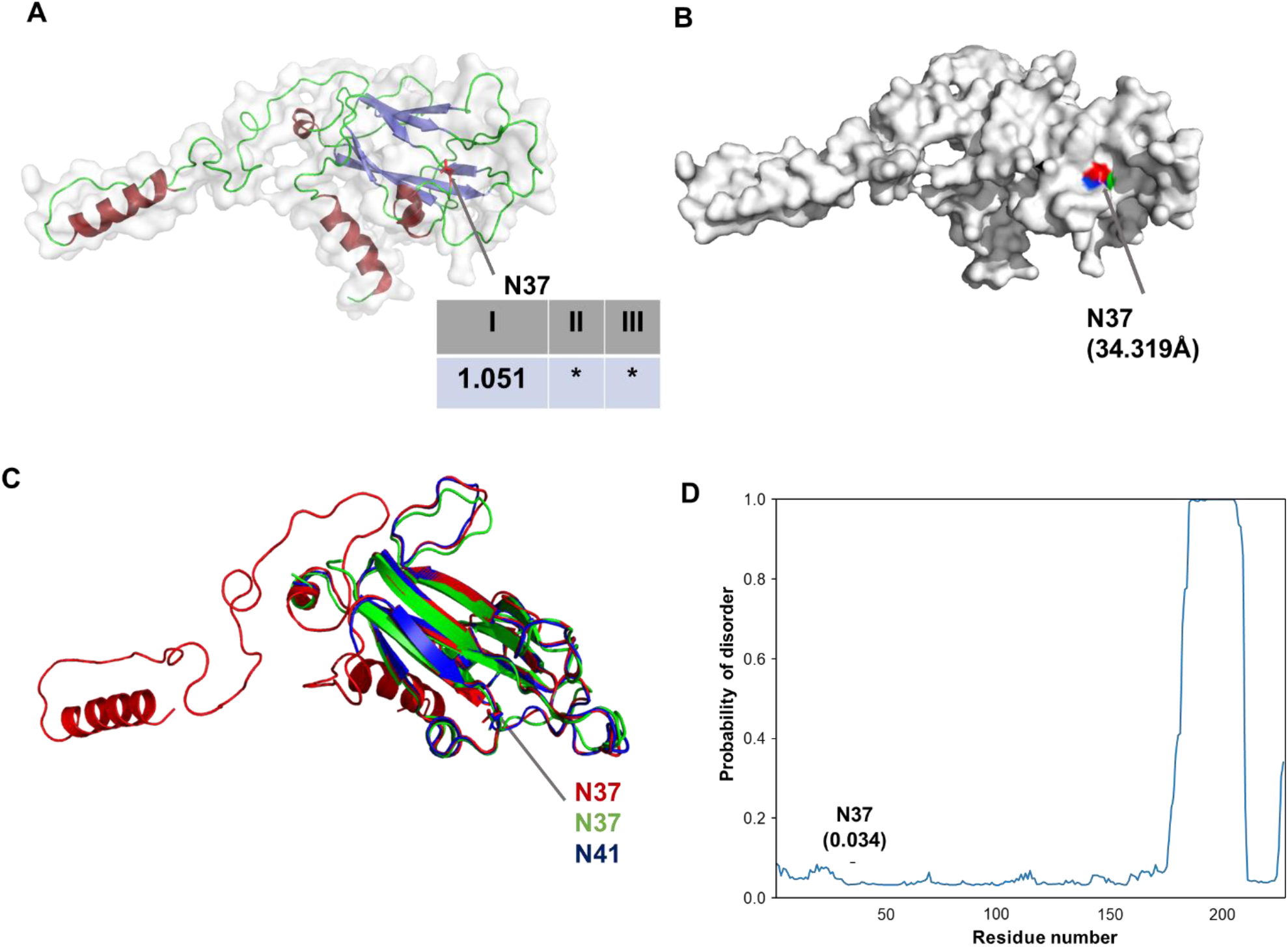
3D structure of ephrin-A5 (F7GZC7) showing the mapped N-glycosylation site. **A**) Cartoon representation of the modelled 3D structure of ephrin-A5 illustrating the ratios of the MS/MS intensities of FNGPs between PNGase F-treated and -untreated cell extracts. α-helix are in firebrick red, β-sheet in slate blue, and loops in green. **I** represents the intensity ratios of CTRL+PNGase F/CTRL, **II** represents the ratio between DTT+PNGase F/DTT, and **III** represents the ratio of **II**/**I**, **\*** represents ratios with DTT+PNGase F equal to 0, indicating deglycopeptides exclusively missing in DTT+PNGase F fractions. **B**) Surface representation showing the SASA of the N-glycosite calculated using PyMol, where the warmer and cooler colors represent N-glycosites with more and less surface exposure, respectively. The SASA of the mapped N-glycosite (in Å) is shown in parenthesis. **C**) 3D structure alignment of modelled *M. mulatta* (red) ephrin-A5 and determined 3D structures of homologs from *H. sapiens* (green) and *M. musculus* (blue) illustrating the conserved localization of the N-glycosite undergoing pronounced structural conformational modulation upon DTT treatment. The N-glycosites are represented using sticks to show the side chains of the glycosylated Asn. **D**) Localization probabilities of ephrin-5A amino acids calculated using ODiNPred, with the mapped N-glycosite (N37) highlighted.

Two of the proteins identified in subset iii) included proteins involved in the protein glycosylation machinery; hexosyltransferase (F6UCJ0/ B3GALNT1) and polypeptide N-acetylgalactosaminyltransferase (A0A1D5QY41/GALNT10). These membrane enzymes reside in the Golgi apparatus. GALNT10 transfers N-acetylgalactosamine from UDP-N-acetyl-α-D-galactosamine to proteins in the Golgi. This enzyme has 6 putative N-glycosites and 13 cysteines. Three N-glycosylation sites have been mapped on the human homolog of N-acetylgalactosaminyltransferase (Q86SR1/ GALNT10) at positions N124, N146 and N593 [95], glycosites also conserved in *M. mulatta* GALNT10. In addition, the human homolog has 5 disulphide bridges at positions Cys135 ↔ Cys365, Cys356 ↔ Cys432, Cys471 ↔ Cys488, Cys523 ↔ Cys538 and Cys563 ↔ Cys578 [95] sites that are also conserved in *M. mulatta* GALNT10, suggesting close structural similarity between human and *M. mulatta* GALNT10 based on primary amino acid sequence similarities. In our analysis, we identified two FNGPs at positions N128 and N575 (**Figure 5A**), corresponding to N146 and N593, respectively, in the human homolog. At position N128, the N-glycosite was determined to be exclusively missing in the DTT and PNGase F treated fraction, suggesting a pronounced structural modulation at this glycosite compared to the control fractions. Three-dimensional analysis of the modelled protein structure showed that the N-glycosite at this position is located on a loop-alpha helix interface (**Figure 5A**). The N-glycosite at N575 showed that this site underwent subtle structural changes, allowing PNGase F to cleave N-glycans at a higher rate upon DTT treatment compared to control fractions at a ratio of 0.48 (**Figure 5A**). This site is also located on a loop proximal to a beta sheet (**Figure 5A**). The SASA of N128 and N575 were 60.4 Å and 47.5 Å, respectively (**Figure 5B**), illustrating that under native conditions, N575 is more occluded compared to N128, corroborated by limited deglycosylation assay (**i**) (**Figure 5A**).

**Figure 5.**
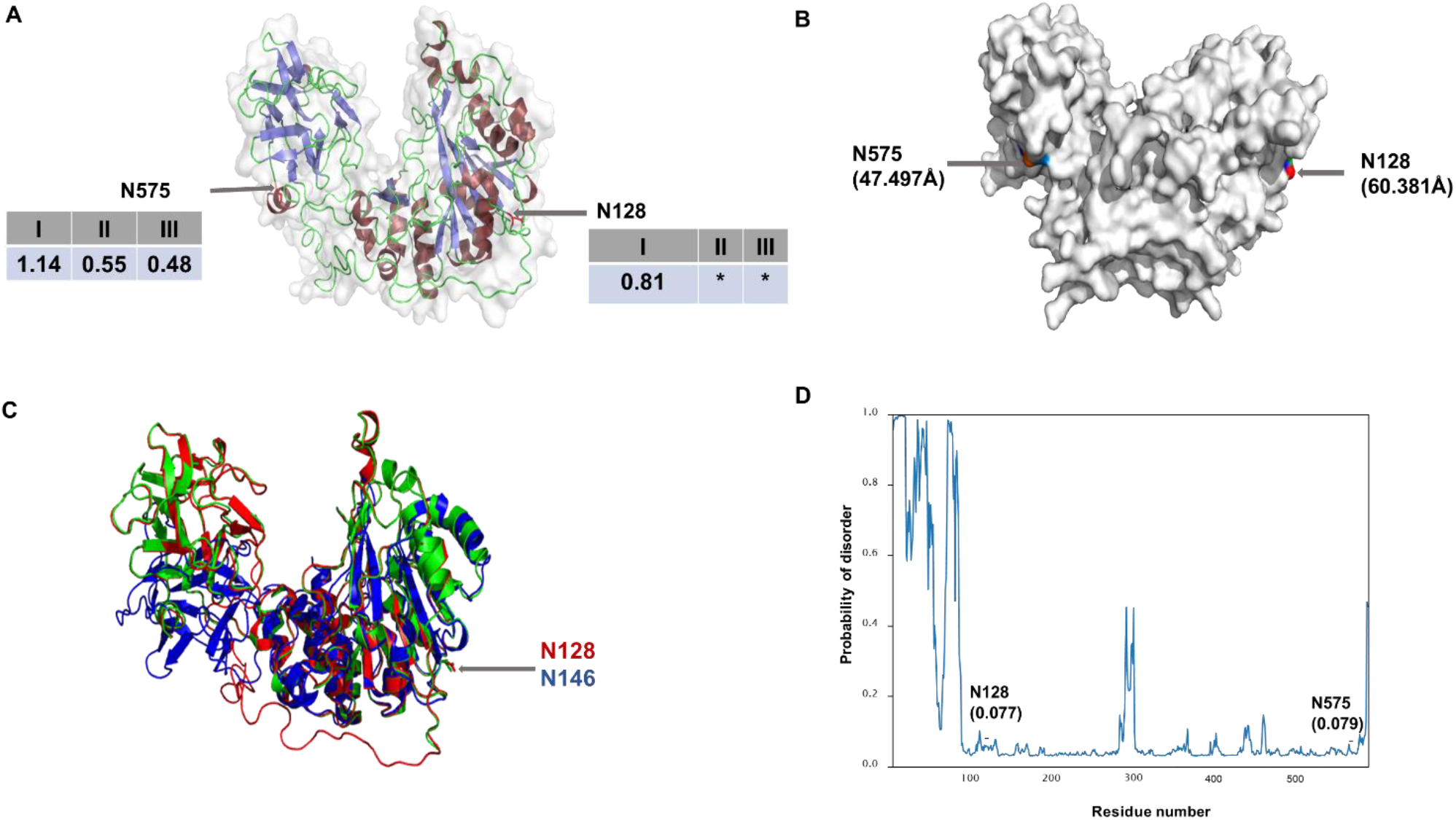
3D structure of GALNT10 showing N128 and N575 N-glycosylation sites undergoing structural modulation upon DTT treatment. **A**) Cartoon representation of modelled 3D structure of *M. mulatta* N-acetylgalactosaminyltransferase illustrating the ratios of the intensities of FNGPs upon LDA; α-helix are in firebrick red, β-sheet in slate blue, and loops in green. **I** represents the intensity ratios of CTRL+PNGase F/CTRL, **II** represents the ratio between DTT+PNGase F/DTT, **III** represents the ratio of **II**/**I**, **\*** represents ratios whose DTT+PNGase F/DTT is 0, to denote FNGPs exclusively missing in DTT+PNGase F fractions. **B**) Surface representation showing the SASA of the N-glycosites calculated using PyMol. **C**) 3D structure alignment of modelled *M. mulatta* (red) GALNT10 and determined 3D structures of homologs from *H. sapiens* (green) and *M. musculus* (blue) illustrating the conserved localization of the N-glycosite undergoing pronounced structural conformational modulation upon DTT treatment. The N-glycosites are represented using sticks to show the side chains of the glycosylated Asn. **D**) Localization probabilities of the GALNT amino acids calculated using ODiNPred are shown. The probabilities for the N-glycosites to fall into disordered regions have been indicated.

The intensity ratio of the deglycopeptides at N128 and N575 between CTRL + PNGase F and CTRL is 0.82 and 1.14, respectively (**Supplementary Table 2**), corroborating the higher accessibility of N128 by PNGase F cleavage under native conditions compared to N575. N575 becomes more accessible to PNGase F after incubation with DTT, as indicated by a relatively high ratio of 0.55 between the DTT + PNGase F and DTT only fractions. The ratio between DTT and control fractions is 0.49, confirming that under DTT conditions, N-glycans at this site are more prone to undergo PNGase F cleavage compared to non-DTT treated. The 3D structures of the human homolog of GALNT10 (2D7I) and *M. musculus* (1XHB) have been determined [95, 96], and their structures were aligned with the modelled *M. mulatta* GALNAT10 glycoprotein, highlighting structural conservation between *M. mulatta* and *H. sapiens* N128 (**Figure 5C**). In *M. musculus*, this N-glycosite is not conserved. Our calculations showed low probabilities (0.077 and 0.079, respectively) for N128 and N575 to localize in disordered regions of the proteins (**Figure 5D**). In addition, the neighbouring amino acids for the two N-glycosites also have low probabilities predictions to be located within disordered regions.

The glycoprotein A0A5F8ABK3 was identified with 4 FNGPs, of which N278 was exclusively missing in the DTT+PNGase F treated fraction. The human homolog of the poliovirus receptor (PVR, P15151/ CD155) is a membrane glycoprotein with three immunoglobulin-like domains in the extracellular N-terminal tail. Human PVR was originally identified as poliovirus receptor, in which the V-type domain is necessary and sufficient for viral binding and internalization [97, 98]. Moreover, PVR binds vitronectin [99]. The human homolog has 10 N-glycosylation sites at N105, N120, N188, N218, N237, N278, N307, N313, N352, and N360. In our analysis, we identified 4 FNGPs at N120, N278, N307, and N313 (**Supplementary Table 2**). 3D analysis of the modelled *M. mulatta* PVR showed that 3 of the 4 N-glycosites were located in loops, with the exception of N120 located on a beta sheet (**Figure 6A**). The N-glycan of the human PVR glycoprotein at N120 was previously reported in human urine samples following MS/MS analysis of intact glycopeptides, and determined to be a complex-type glycan (Hex5HexNAc4) [100]. N120 was more occluded compared N-glycosites within loops under both DTT treated and untreated conditions as shown by the ratios between CTRL+PNGase F/CTRL, DTT+PNGase F/DTT. Our analysis showed that N278 had the lowest SASA value among the sites located on loops (48.4 Å) compared to N307 and N313 sites with 56.4 Å and 63.9 Å, respectively (**Figure 6B**), but N278 were more susceptible to PNGase F cleavage under native conditions (**Figure 6A**). Upon DTT treatment, the N278 was completely deglycosylated, illustrate that this site became completely accessible to cleavage by PNGase. Alignment of the human PVR showed that this site is located on a beta sheet (**Figure 6C**). The glycosylation of N150 and N120 is not essential for viral replication and tissue tropism [101]. Removal of 4 glycosylation sites in the two outermost V and C-like domains did not influence binding to vitronectin [99]. The probabilities of the N-glycosites to fall within disordered regions were determined to be 0.0359, 0.0384, 0.0343 and 0.0329 for N120, N278, N307 and N313, respectively (**Figure 6D**).

**Figure 6.**
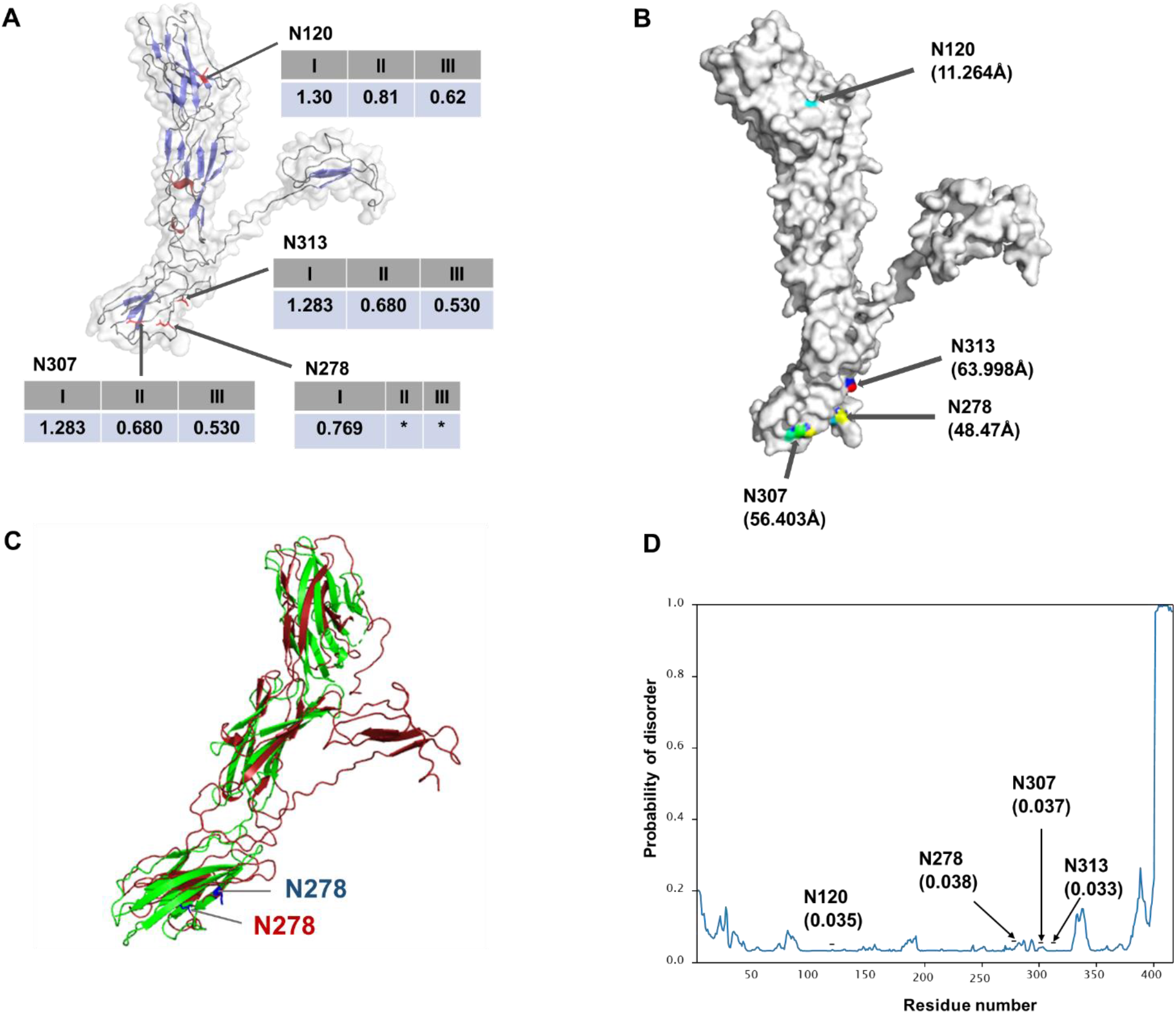
3D structure of the uncharacterized protein (A0A5F8ABK3/PVR) showing N-glycosites of interest. **A**) Cartoon representation of PVR illustrating the ratios of the intensities of FNGPs. α-helix are in red, β-sheet in blue, and loops in grey. The N-glycosites are in red, and sticks used to represent the side chains. **I** represents the intensity ratios of CTRL+PNGase F/CTRL, **II** represents the ratio between DTT+PNGase F/DTT, and **III** represents the ratio of **II**/**I**, * represents ratios whose DTT+PNGase F is 0, representing deglycopeptides exclusively missing in DTT + PNGase F fractions. **B**) Surface representation showing the SASA of the N-glycosite calculated using PyMol, where the warmer and cooler colors represent sites with more and less exposure, respectively. The SASA (in Å) are shown in parenthesis. **C**) Structure alignment of the modelled *M. mulatta* PVR and the determined structure of the human homolog (QARQ) deposited in PDB. N278 is indicated on both proteins. **D**) Localization probabilities of the PVR amino acids calculated using ODiNPred. N-glycosites of interest are highlighted.

FNGPs were also identified and quantified in our analysis showing no changes in N-glycosite exposure (and by inference no structural changes in those regions) between CTRL and DTT treated fractions. An example, the basal cell adhesion molecule (A0A1D5Q3L5/BCAM) which was identified with 4 FNGPs; including 2 FNGPs located on beta sheets (N299 and N417), and 2 FNGPs located on loops (N355 and N361) (**Supplementary Figure 4**). The human homolog has been mapped with 5 putative N-glycosites (N321, N377, N383, N419, N439). Human BCAM is known to be involved in adhesion of cells and extracellular matrix components such as laminin, and has been shown to be a receptor of laminin in malignant tumors [102]. The 3D-structure of the modelled *M. mulatta* BCAM revealed that N-glycosites not affected by DTT treatment are located in beta sheets (**Supplementary Figure 5A**), as indicated by their low SASA (67.8 Å and 34.4 Å, respectively) (**Supplementary Figure 5B**). N355 and N361 glycosites were affected by DTT treatment (**Supplementary Figure 5A**). These sites are located in loops as indicated by structural modelling (**Supplementary Figure 5A**), and analysis of their localization probabilities indicate that they are located in areas with low probabilities of disorder (**Supplementary Figure 5C**).

The presence of specific PTMs in structural defined motifs has been shown for phosphorylated residues located within unstructured disordered regions both in eukaryotes and prokaryotes [103]. The phosphorylation of these disordered regions allows for enhanced and stabilized protein-ligand binding [104]. Protein O-glycosylation has been mapped also to intrinsically disordered regions protecting specific domains against proteolysis [105, 106]. N-glycosylation has been shown to impact glycoprotein 3D structure and folding through the ER-resident quality control system and thereby is directly involved in dictating the fate of glycoproteins [107, 108]. Thermodynamic and experimental analyses have shown that N-glycosylation often stabilize the glycoprotein structure [109, 110]. N-glycosites are found mainly in ordered regions; amino acid residues surrounding N-sequons are relatively poor in hydrophobic and bulky residues known to promote disordered regions [94]. Proteome-wide structural analysis of N-glycosites from human, mouse, fly, plant and yeast showed that a high proportion of N-glycosites are localized within a turn/loop and are solvent-exposed [111–113]. Approximately 80% of all N-glycosites of the human proteome are located in loop/turn and this pattern is consistent in other eukaryotes [113]. Our data shows that the highly exposed N-glycosites located in turn/loops are more prone to undergo conformational changes upon stress induction compared to sites within alpha helix or beta sheet structures. Moreover, our analysis confirms previous findings showing N-glycosites located in ordered regions. The presence, abundance and occupancy of N-glycosylation enriches the structural features of a glycoprotein that cannot be derived solely based on the primary protein sequence thus conferring a large diversity of functions. The applicability of this method to studying changes in glycoprotein conformation during stress induction can readily be used to study glycoprotein structural changes under other physiological and pathological conditions.

## Conclusion

Here we describe a new systems-wide method termed LDA that probes the structural changes of glycoproteins under controlled conditions. LDA explores the differential accessibility of PNGase F to N-glycosites depending on the folding state of a glycoprotein and uses MS-based proteomics to quantify altered abundance in FNGPs. We developed and validated the method using standard glycoproteins and then showed the potential of the method by evaluating the conformational changes of complex mixtures of glycoproteins in LLC-MK2 cells treated with DTT. Strikingly, the N-glycosites undergoing the highest conformational changes under ER stress induction are located on loops, corroborating findings from previous studies using limited proteolysis. LDA can be applied to study the glyco-proteostasis of different cells or tissues in health and disease, to explore the system-wide modulation of glycoprotein structures in a complex cellular milieu. Since this method is based on MS readout, the detection of most abundant glycoproteins is favoured and an average conformation is detected when multiple glycoprotein folding states co-exist. Moreover, the different accessibility of PNGase F to a specific glycosylation site could be due to differential site occupancy, aggregation, protein-protein interaction, small molecule binding and/or conformational changes. These aspects can be seen as a limitation since it requires specific information on the origin of the changes and a strength since it can be applied to several conditions. The introduction of intact glycopeptide analysis, different enrichment methods, advanced MS-based proteomics tools and combination with other structural approaches will enhance the depth of this method and is thus a logical next step in further method development. We believe that the presented LDA method will serve as a platform to develop other limited enzymatic assays to probe conformational changes based on PTM analysis. Overall, the glycoprotein conformational features deduced from the LDA correlate with those deriving from other biophysical and spectroscopic techniques. LDA will thus contribute to the diverse toolbox required to better study glycoprotein structure and function.

---

**Supplementary Figure 1.**
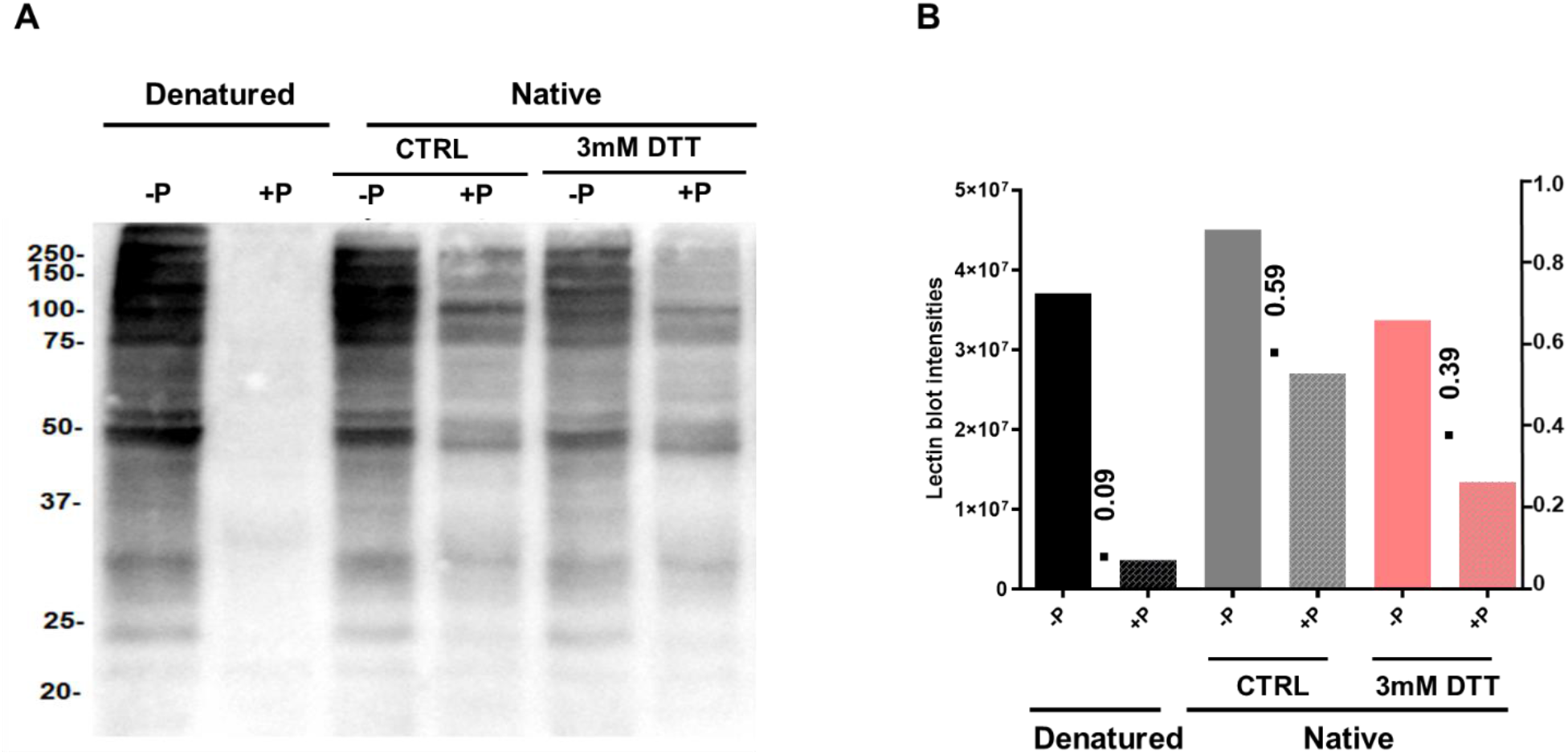
Optimized LDA applied to LLC-MK2 cellular glycoproteins. **A**) Lectin blotting using Con A on LLC-MK2 cells following LDA using optimized conditions previously determined on standard glycoproteins. **B**) Intensity and ratios between +PNGase F/−PNGase F calculated using ImageLab are shown. - P = negative control not treated with PNGase F; +P = PNGase F treated.

**Supplementary Figure 2.**
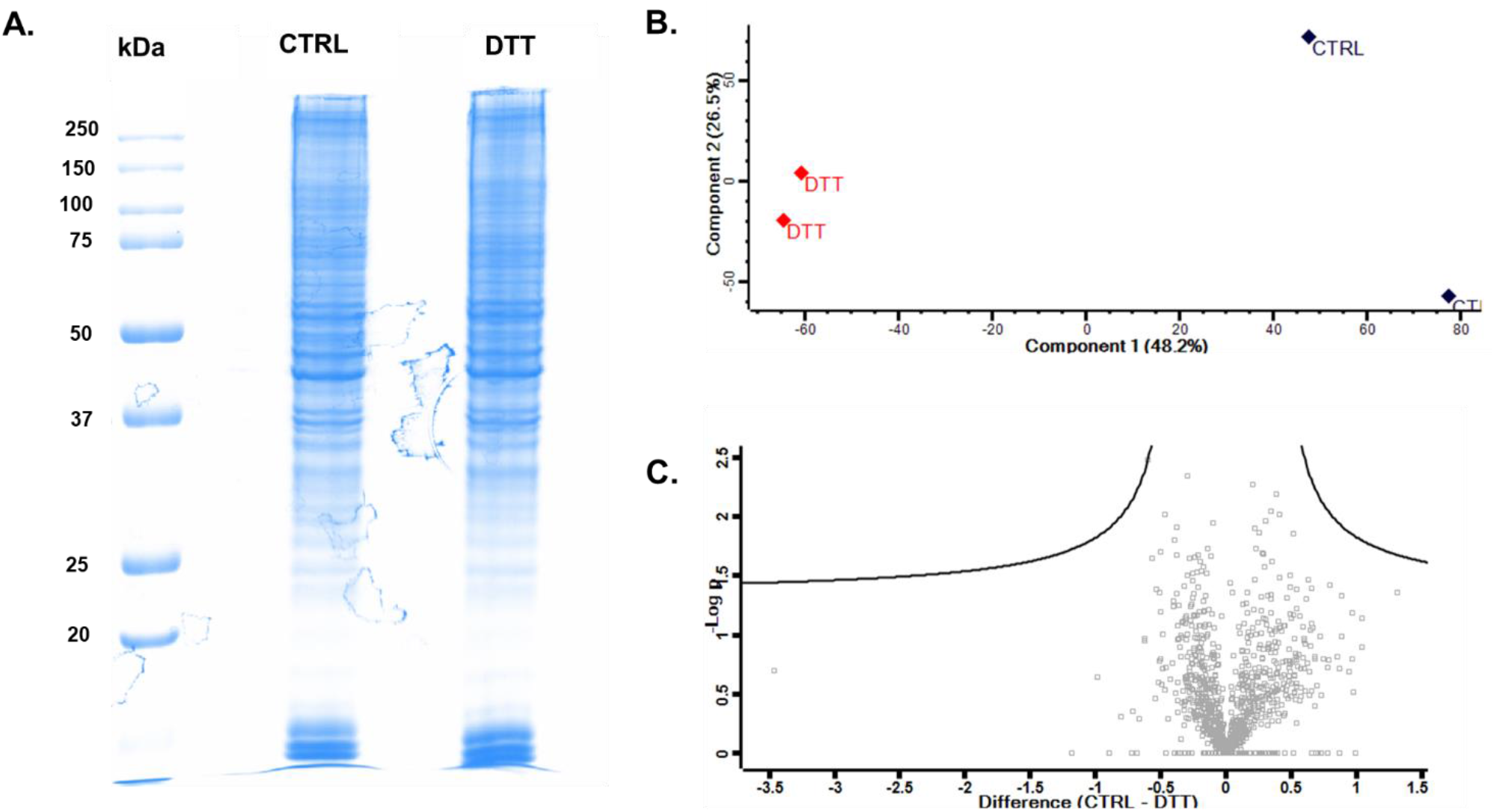
Protein expression analysis of LLC-MK2 cells incubated in medium supplemented with or without 3 mM DTT for 30 min. (**A**) The protein expression profiles were visualized by SDS-PAGE, and the band intensities analyzed using ImageLab. (**B**) The effect of DTT on the proteome was visualized using PCA, and the levels of regulation were visualized using (**C**) Volcano plot calculated at a FDR < 0.05.

**Supplementary Figure 3.**
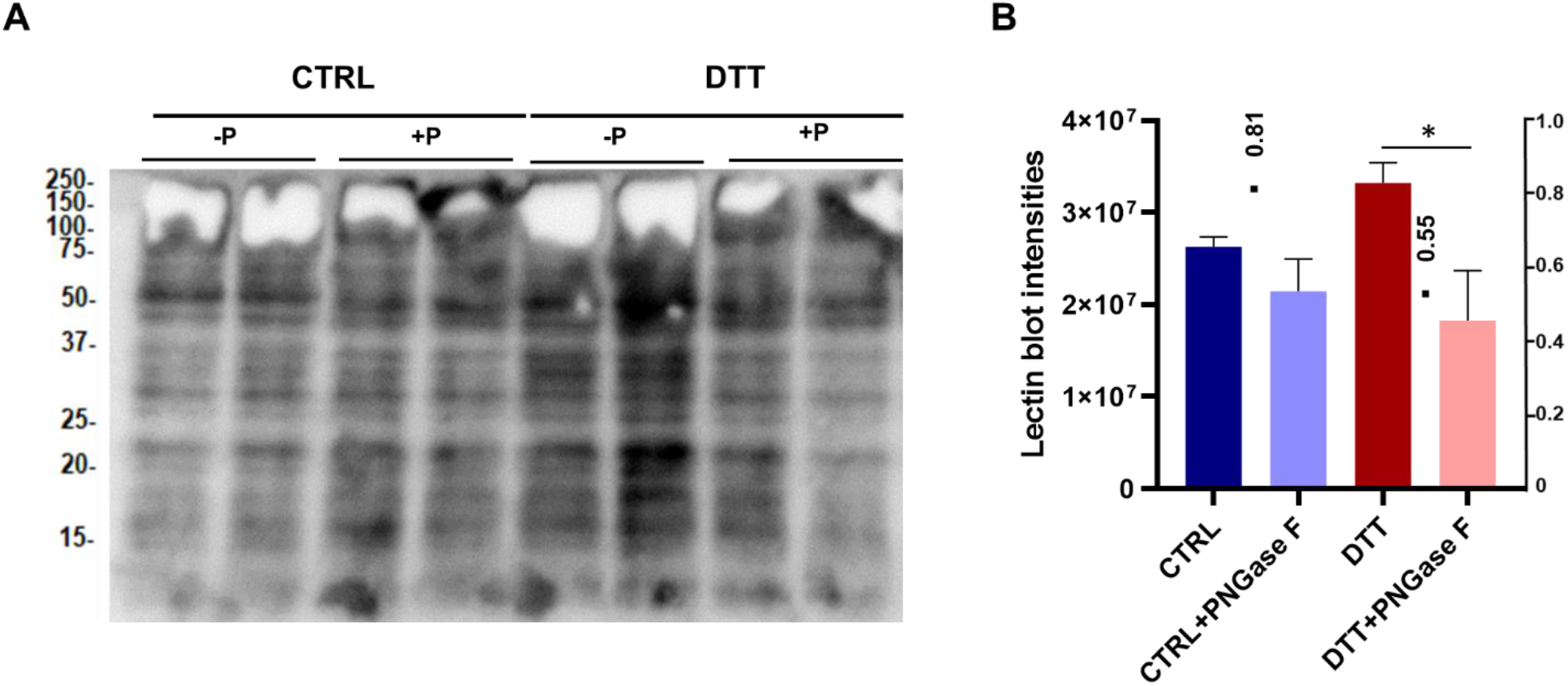
Lectin blot analysis of LLC-MK2 cells incubated with or without 3 mM DTT for 30 min followed by deglycosylation under native conditions. **A)** ConA lectin blot following LDA. **B**) The combined intensities for each condition are summarized, and the ratios between +PNGase F/−PNGase F for both control and DTT treated fractions. Results are displayed as mean ± SEM; *p < 0.05.

**Supplementary Figure 4.**
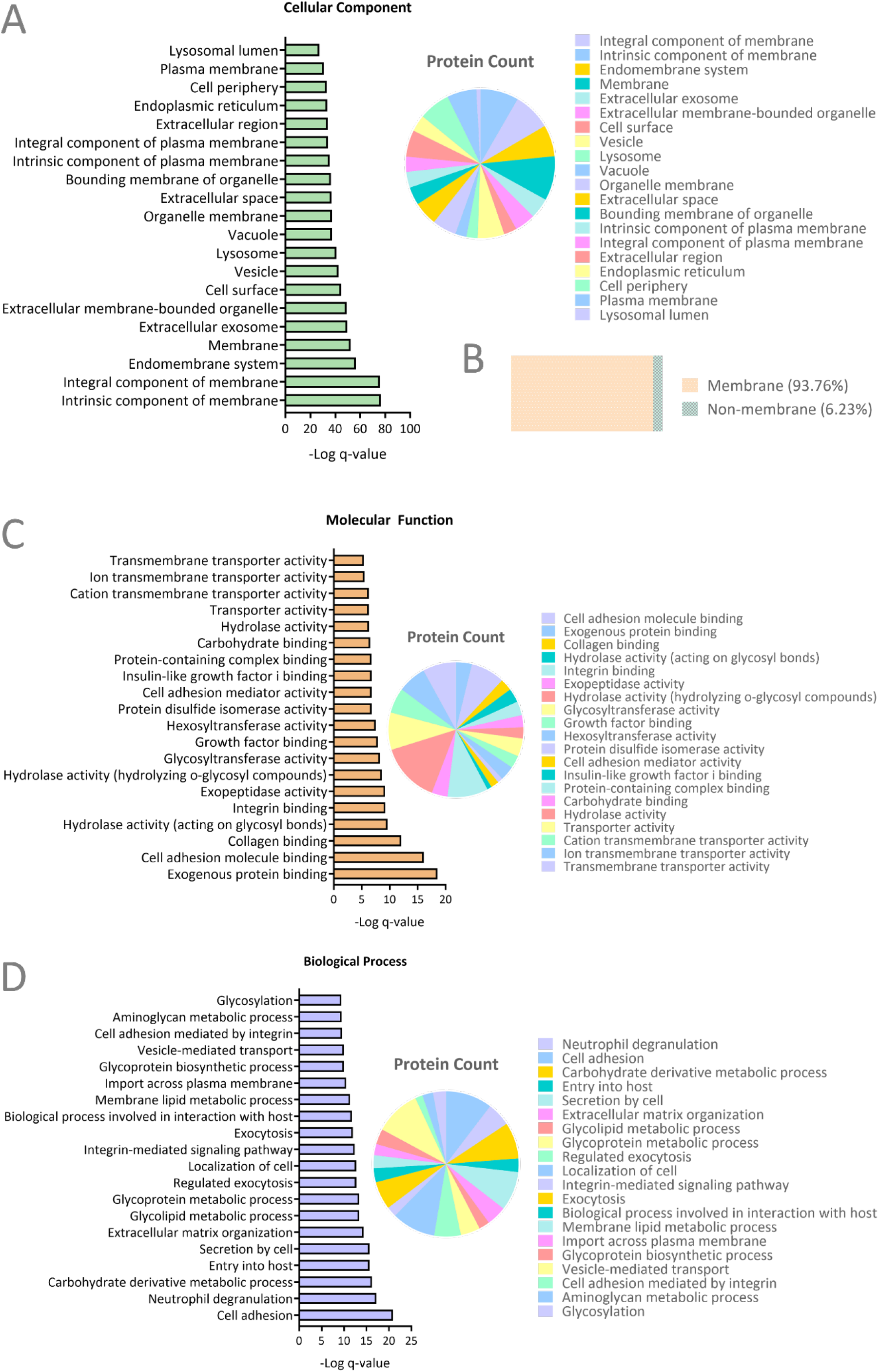
Gene ontology analysis of LLC-MK2 deglycoproteome following LDA. Top 20 Gene Ontology terms of **A**) Cellular components, **B)** percentage of proteins that localize to the membrane. **C**) Molecular functions and **D**) Biological processes are illustrated. Only terms with q-value <0.05 were considered (Benjamini-Hochberg).

**Supplementary Figure 5.**
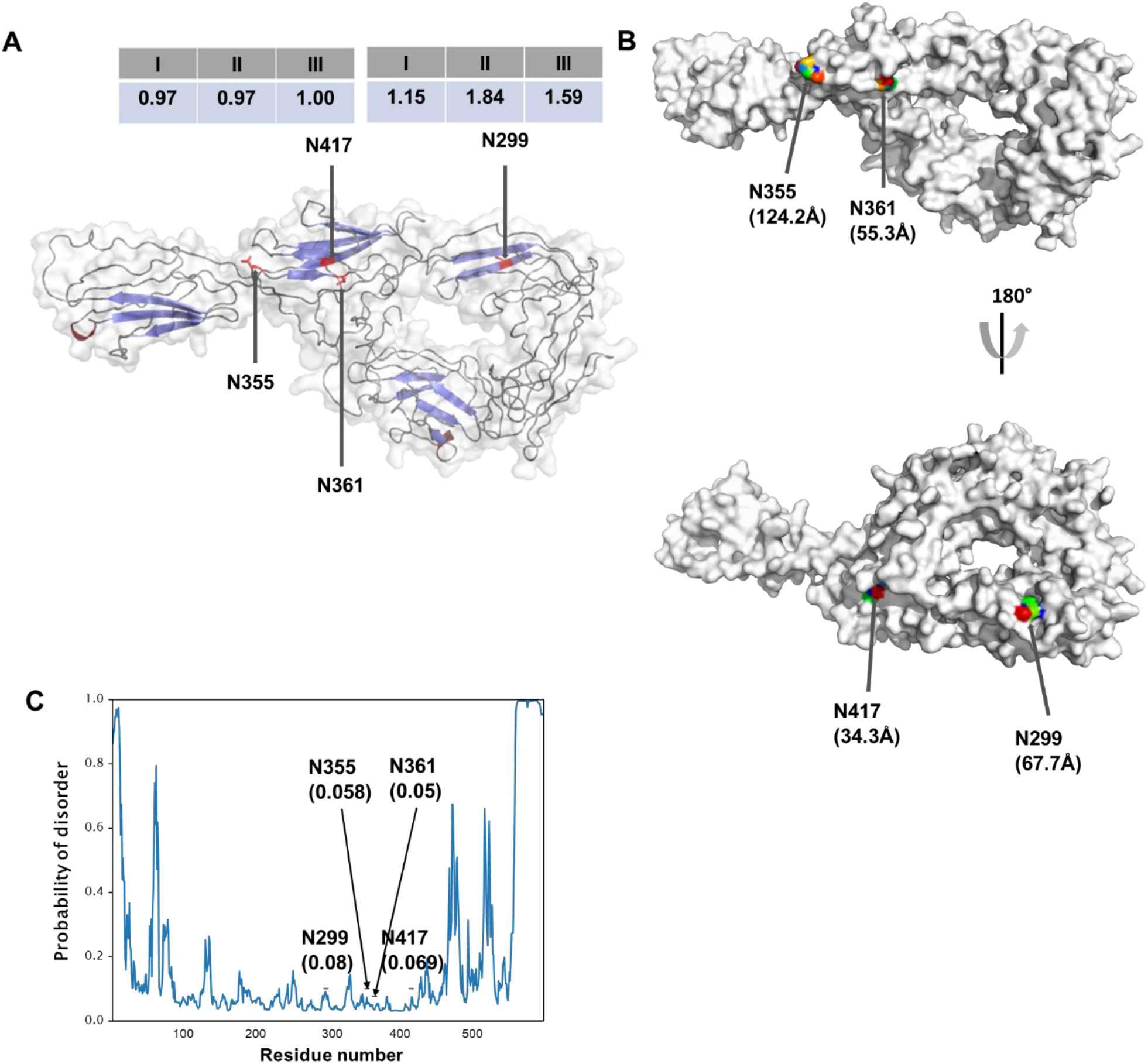
3D structure of basal cell adhesion molecule (Lutheran blood group) (A0A1D5Q3L5/BCAM) showing identified N-glycosites. **A**) Cartoon representation of PVR illustrating the ratios of the intensities of FNGPs. α-helix are in red, β-sheet in blue, and loops in grey. The N-glycosites are represented in red, and sticks represent the side chains. **I** represents the intensity ratios of CTRL+PNGase F/CTRL, **II** represents the ratio between DTT+PNGase F/DTT, and **III** represents the ratio of **II**/**I**, * represents ratios whose DTT+PNGase F is 0, representing deglycopeptides exclusively missing in DTT + PNGase F fractions. **B**) Surface representation showing the SASA of the N-glycosite calculated using PyMol, where the warmer and cooler colors represent sites with more and less exposure, respectively. The SASA (in Å) are shown in parenthesis. **C**) Localization probabilities of the BCAM amino acids calculated using ODiNPred. The N-glycosites of interest are highlighted.

## Conflict of interest

The authors declare no conflict of interest.

## Acknowledgments

Prof. Maria Julia Manso Alves, Prof. Walter Colli and Célia Aparecida Ludio Braga are acknowledged for the LLC-MK2 cells and for the advices during the manuscript preparation. The work was supported by grants and fellowships from FAPESP (2018/18257-1, 2018/15549-1, 2020/04923-0 to GP, 2013/07937-8 to LL, VDM (2021/00399-7) and SNM (2017/04032-5), CNPq (DOQ and “Productivity fellowship” to GP), and CAPES to LRF and JMDS.

## Notes

### Competing Interest Statement

The authors have declared no competing interest.

